# Evolution of the division of labor between two interacting public tasks

**DOI:** 10.64898/2026.10.13.758955

**Authors:** Aidan Fielding, Joshua B. Plotkin, Erol Akçay

## Abstract

Individuals in many species perform several costly tasks whose benefits are shared with neighbors: bacteria secrete distinct public metabolites, plants emit distinct defensive volatiles, and human foragers acquire distinct shared foods. Performing one of these tasks often changes the benefits and costs of the other: a forager who brings in one kind of food provides different nutritional benefits to neighbors than one who brings in several kinds, and the difference depends on what the neighbors bring. Here we ask which divisions of labor evolve when two tasks interact. We model the interaction at both the group and the individual level, and we derive the long-run outcomes of the replicator dynamics as functions of the task payoffs and the interaction parameters. Interactions between tasks produce long-run outcomes that cannot occur when the tasks are independent. In particular, specialists, who perform one task but not the other, can coexist in a stable division of labor only if the tasks interact, and in most cases only if the interaction is more favorable when different individuals perform the two tasks than when one individual performs both. We also show that a specialized division of labor can be bistable with a non-specialized one, and we prove that, when both tasks are identical anti-coordination games, individuals in the specialized population have higher fitness. This bistability produces hysteresis in both the division of labor and fitness as interaction parameters vary. Populations with identical payoffs can therefore evolve to divisions of labor with different mean fitness.

**Non-specialist summary:** Many organisms perform several costly tasks whose products are shared with neighbors: bacteria secrete molecules used by other cells, and human foragers share the different foods they collect. The value of one task often depends on whether the other is also performed, and on whether one individual performs both or two perform one each. We model evolution mathematically to ask how such interactions determine who performs which task. Individuals that each perform one task coexist only when the tasks interact, and usually only when two individuals who split the tasks gain more from their interaction than one individual who performs both. A population can evolve to either of two divisions of labor with different mean fitness, depending on its initial composition.

## 1 Introduction

Individuals in many species can perform more than one costly task whose benefits are shared with neighbors. Each task is a separate act of cooperation: it consumes the performer’s own time, energy or metabolic capacity, and it yields a good that others also consume and benefit from. Among bacteria, for example, species produce a variety of publicly beneficial extracellular metabolites, sometimes producing two or more within a colony (Drescher et al., 2014; Kramer et al., 2020; Smith and Schuster, 2019). A well-studied example is *Pseudomonas aeruginosa*, which produces two different types of iron-scavenging molecules called siderophores (Ross-Gillespie et al., 2015). In *Bacillus subtilis* colonies, cells can differentiate to produce surfactin or extracellular matrix, both of which are needed for colony expansion (van Gestel et al., 2015). Bacteria also engage in bidirectional cross-feeding, in which two cell types each produce a metabolite used by the other (Özkaya et al., 2017). The social amoeba *Dictyostelium discoideum* performs two tasks, fruiting body formation and macrocyst formation, that both aid in survival during starvation (Shibasaki and Shimada, 2018). Turning to multi-cellular organisms, the mound building mouse *Mus spicilegus* may transport either vegetation or roof-building materials during nest construction, both of which are essential for the completion of the nest (Hurtado et al., 2013; Serra et al., 2012). Plants emit a range of volatile compounds, each a separate metabolic task, that defend them and their neighbors from insect herbivores (Creagar et al., 2025). In human hunter-gatherers, what each individual collects is often shared, so foraging for each type of shared food is a distinct task with a public benefit and a private cost (Anderson et al., 2023; Bird, 1999; Stibbard-Hawkes et al., 2022).

How these tasks are divided among the individuals of a population, that is, who performs which task, if any, is then an important question. A population may be monomorphic, with everyone taking the same action (perform or not) on a given task, or polymorphic, in which case several distinct arrangements are possible. Specialization is the arrangement in which two types coexist, each performing one task and leaving the other to its partner. An alternative division of labor is one type performing both tasks and another performing neither. The population may also contain a mixture of these arrangements. Which of these arrangements evolves, and what each implies for the individuals in it, is the subject of this article.

A large body of theory asks when specialists are favored over generalists, and these studies share a common requirement: an individual that concentrates on one task must obtain more from that task than an individual that divides its effort between two. Rueffler and colleagues showed that division of labor between two functions is favored when the performance functions accelerate, together with positional effects and synergistic interactions among the parts that perform them (Rueffler et al., 2012). Cooper and West obtained the same requirement for a cooperative behavior, and showed that a sufficiently large efficiency benefit selects for extreme specialization, in which some individuals become wholly dependent on the helping of others (Cooper et al., 2021; Cooper and West, 2018). Related arguments underlie accounts of the origin of multicellularity (Gavrilets, 2010; Ispolatov et al., 2012; Michod, 2006; Yanni et al., 2020), of metabolic division of labor among microbes (Giri et al., 2019; Tsoi et al., 2018; West and Cooper, 2016), of task allocation in social insects (Beshers and Fewell, 2001; Duarte et al., 2011), and of returns to skill in human foraging (Hooper et al., 2015). In every one of these accounts, the benefit of specializing is a property of how well a single individual performs a single task.

In the simplest theoretical model of this situation, each task corresponds to a two-person, two-strategy game in which players can have the pure strategies *C* (perform task; cooperate) or *D* (not perform task; defect). Individuals play two of these task-games simultaneously with a coplayer drawn uniformly at random from the population, and their fitness is proportional to the expected payoff they derive from both tasks. The standard assumption is that the tasks are independent, so that the payoff from both tasks is the sum of the two single-task payoffs. Cressman et al. (2000) and Hashimoto (2014) showed that replicator-dynamical equilibria in the two-game, two-strategy model can be directly inferred from the equilibria of the single games, analyzed separately. Of the three canonical regimes of a single game, only the anti-coordination game has a polymorphic equilibrium, at which both strategies coexist; therefore, in the model of two independent games, for two or more strategic types to coexist, at least one of the games must be an anti-coordination game. When both games are anti-coordination games, there is a line of evolutionary fixed points toward which the population evolves. Cressman et al. (2000) showed that the rest point on this line for a given evolutionary trajectory is determined by the initial strategy frequencies via an invariant manifold. Hashimoto (2014) showed that a finite population in this regime evolves to the line of rest points, and stochastic perturbations move the population along it. Chamberland and Cressman (2000), Hashimoto (2006), and Venkateswaran and Gokhale (2019) demonstrated further that when games have more than two strategies, equilibria in the two-game model cannot be inferred from those of the independent games.

In nature, however, performing one task often changes the payoff of the other. In *P. aeruginosa* colonies, the presence of one type of siderophore (pyoverdine) suppresses production of the other type (pyochelin) (Ross-Gillespie et al., 2015), and clones that stop producing siderophore also form less biofilm (Harrison and Buckling, 2009). In *B. subtilis* colonies, secreted surfactin reduces friction between the cells and their substrate, which facilitates construction of the extracellular matrix and overall colony growth. Similarly in the amoeba *D. discoideum*, fruiting body and macrocyst formation behaviors are controlled by the same chemical signal, making the performance level of one task dependent on that of the other. For the mound building mice, the benefits of collecting vegetation and roof materials completely depend on each other, because both kinds of material are required for a functional nest. Among humans, foraging different foods may have complementary benefits if they are from different nutritional categories, substitutable benefits if they are from the same category, or antagonistic benefits if they have an undesirable chemical reaction that reduces nutritional quality. Furthermore, an individual who forages one food may pay an opportunity cost because they have less time or energy to forage another. They may also benefit by transferring skills used to forage one food to foraging another.

The theory of division of labor and the theory of two simultaneous games therefore leave a question open between them. Efficiency arguments for specialization concern how well one individual performs one task; the theory of two simultaneous games concerns two tasks that do not affect one another. Neither describes the examples above, in which performing one task changes the payoff of the other. Three questions motivate our analysis: Which divisions of labor evolve when two tasks interact? Does complementarity between tasks favor specialists, as intuition suggests? And what does the resulting division of labor imply for the fitness of the individuals living in it?

Here we develop a model of the evolution of two interacting tasks, by adding to the standard model of two simultaneous games a term for the change in payoff when both tasks are performed. We allow that change to differ between individuals who perform both tasks and individuals who perform at most one, which is how a resource constraint or a transfer of skill between tasks enters the payoffs. When the tasks are anti-coordination games with identical benefits and costs, interactions produce twelve distinct evolutionary outcomes. Six of these outcomes are monostable; in the other six, two or three divisions of labor are stable, and the initial composition of the population determines which one evolves. A division of labor containing both specialists is the unique stable equilibrium in four outcomes, which occur with either complementarity or antagonism between the tasks. In three further outcomes specialization is bistable with an asymmetric division of labor in which one type performs both tasks and the other performs neither, and these three outcomes require every type to benefit from complementarity between the tasks. In five of these seven outcomes, specialists persist only if the interaction is more favorable for specialists and non-performers than for performers of both tasks, a condition analogous to the efficiency requirement of the division-of-labor literature. We also show that, when both tasks are anti-coordination games, any interaction between them, however small, replaces the line of rest points of the additive model with isolated rest points. We prove analytically that individuals in the specialized population have higher fitness in every case of bistability between a specialized and an asymmetric division of labor. We also show that bistability produces hysteresis in both the division of labor and mean fitness when the level of complementarity between the tasks oscillates.

## 2 A model of two interacting tasks

Each task is a two-person, two-strategy symmetric game with pure strategies “perform task” (*C* ) and “not perform task” (*D* ), and payoffs to the row player given by

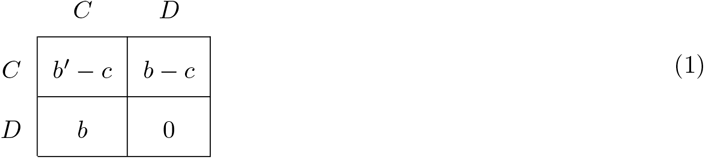

In matrix 1, *c* is the private cost of performing the task, *b* is the per-capita benefit when one player performs the task, and *b*^*′*^ *> b* is the per-capita benefit when both players perform the task. The benefit of a task is public: a player who does not perform the task still collects the benefit *b* produced by a coplayer who does. The task with payoffs defined by matrix 1 has a polymorphic equilibrium, with cooperators and defectors coexisting, if and only if it is an anti-coordination game, which is defined by the inequalities *b − c >* 0 and *b*^*′*^ *− c < b*. The first inequality states that performing the task pays when nobody else does, and the second inequality states that performing it does not pay when the coplayer already has: returns to the task diminish. An anti-coordination game with this payoff structure is also called a Snowdrift game, and it is the standard model of a public good whose producers and non-producers coexist (Doebeli and Hauert, 2005; Doebeli et al., 2004; Gore et al., 2009).

An individual’s fitness *f* is given by the expected payoff it will receive if its coplayer is drawn uniformly randomly from the population. From matrix (1), denoting the frequency of individuals with strategy *C* (*D*) in the population as *x*_*C*_ (1 *− x*_*C*_), the fitness of *C* individuals is *f*_*C*_ = *x*_*C*_(*b*^*′*^ *− c*) + (1 *− x*_*C*_)(*b − c*) and that of *D* individuals is *f*_*D*_ = *x*_*C*_*b*. The evolution of *x*_*C*_ over generational time is given by the replicator equation

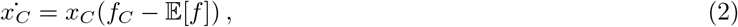

where E[*f*] is the expected fitness in the whole population. If the task is an anti-coordination game, there is an interior, evolutionarily stable frequency 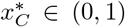 of task performers, so *C* and *D* individuals coexist in the population with frequencies 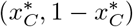.

In our setting, two tasks affect fitness. The set of possible strategies is (*CC, CD, DC, DD*), where the first (second) letter of each strategy indicates its action in task one (two). An individual with strategy *CC* performs both tasks, and we call such individuals double-cooperators; an individual with strategy *DD* performs neither, and we call such individuals double-defectors. The two remaining strategies, *CD* and *DC*, each perform one task and leave the other to a coplayer, and we call them specialists in task one and task two respectively. Following Cressman et al. (2000) we call the double-cooperators and double-defectors together the generalists, since neither distinguishes between the two tasks. In the standard model of two simultaneous games the total payoff is the sum of the two single-task payoffs (Cressman et al., 2000; Hashimoto, 2006), which corresponds to *δ* = *γ* = 0 in matrix (3) below.

We incorporate complementary or antagonistic interactions between tasks by modifying this additive payoff matrix such that if both tasks are performed, then the total payoff from them changes. We need to distinguish between public and private interactions between tasks. In the former case, the interaction affects both players, e.g., when it happens after the products of performing the tasks are shared. The effect of the public task interaction on a focal individual depends on the actions taken by its coplayer: a task one (two) specialist will only experience the interaction effect if their coplayer is a task two (one) specialist or a double-cooperator, and a double-defector will only experience it if their coplayer is a double-cooperator. We denote this public interaction effect by *δ*; it accrues when both tasks are performed in the group, but only to those who perform at most one task (*CD, DC*, and *DD*). If the same individual performs both tasks, they are also potentially subject to the private interaction between tasks (e.g., the additional cost of having to invest in both tasks, or any interaction between the products of the tasks that can happen *before* the products are shared). The interaction term for double-cooperators can therefore differ from the public interaction term. We denote this interaction by *γ*. A double-cooperator covers both tasks by itself, so it receives its interaction term *γ* against every coplayer; a specialist or a double-defector receives its interaction term *δ* only when a coplayer supplies the task(s) it does not perform. We call an interaction between tasks *complementary*, or equivalently synergistic, when performing both yields more than the sum of the two separately, and *antagonistic* when it yields less. When the two interaction terms differ in sign, different strategies experience complementarity and antagonism at once. The difference between the two interaction terms, *γ − δ*, is what an individual gains or loses by covering both tasks itself rather than leaving one to a coplayer. A negative difference, *γ < δ*, can represent a resource constraint: an individual that divides a fixed budget of time or metabolic capacity between two tasks obtains less from their interaction than one that concentrates on a single task. A positive difference, *γ > δ*, can represent a transfer of skill or machinery between the two tasks, which is available only to an individual that performs both (the difference *γ − δ* can also be regarded as the net result of these two private effects).

Denote by *c*_*k*_ the private cost of performing task *k*, by *b*_*k*_ the per-capita benefit when one player performs task *k*, and by 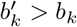 the per-capita benefit when both players perform task *k*, where *k* ∈ {1, 2}. The difference 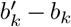 is the more commonly studied interaction between the contributions of two players to the *same* task. The payoff matrix resulting from our assumptions is given by (3), and Table 1 collects every parameter and variable of the model.

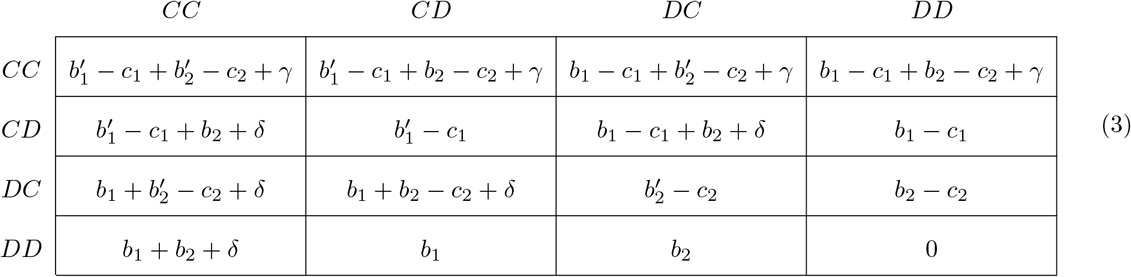

**Table 1:** Parameters and variables of the model. The two inequalities on *b*_*k*_ and 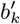 together state that task *k* is an anti-coordination game: performing it pays when the coplayer does not, and does not pay when the coplayer already has. The subscript *k* ∈ {1, 2} indexes the two tasks.

| Symbol | Interpretation | Range |
| --- | --- | --- |
| $b_k$ | per-capita benefit of task $k$ when one player performs it | $b_k > c_k$ |
| $b'_k$ | per-capita benefit of task $k$ when both players perform it | $b_k < b'_k < b_k + c_k$ |
| $c_k$ | private cost of performing task $k$ | $c_k > 0$ |
| $\delta$ | interaction term received by individuals who perform at most one task | $\delta \in \mathbb{R}$ |
| $\gamma$ | interaction term received by individuals who perform both tasks | $\gamma \in \mathbb{R}$ |
| $x_{CC}$ | frequency of double-cooperators, who perform both tasks | $[0, 1]$ |
| $x_{CD}, x_{DC}$ | frequencies of specialists in task one and in task two | $[0, 1]$ |
| $x_{DD}$ | frequency of double-defectors, who perform neither task | $[0, 1]$ |
| $x_C^{*k}$ | frequency of performers of task $k$ when that task is analyzed alone | $(0, 1)$ |

In matrix (3), the interaction term for double-cooperators, *γ*, appears in every entry of the top row and nowhere else, whereas *δ* is absent from the seven entries in which the pair leaves at least one task unperformed: two specialists of the same type, a specialist against a double-defector, and two double-defectors.

The interaction between tasks is realized within an interacting pair rather than through the frequency of each task in the population, and this feature of the model determines the results that follow. The assumption is appropriate when the two products interact locally — within a cell, within a nest, or between two foragers who pool what they collect — and less appropriate for freely diffusing products whose interaction depends on the total amount of each present in the environment. We return to this assumption in the Discussion.

Denote the frequencies of the strategies as (*x*_*CC*_, *x*_*CD*_, *x*_*DC*_, *x*_*DD*_). Throughout we assume a large, well-mixed population in which coplayers are drawn uniformly at random, so that a strategy’s fitness is its expected payoff against the current population. As above, the evolution of the frequency of any strategy, say *CC*, is given by the replicator equation

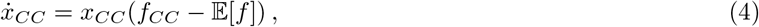

where *f*_*CC*_ is the fitness (expected payoff) of a *CC* individual and E[*f*] is the expected fitness in the current population. Three of the four replicator equations determine the dynamics, since the frequencies sum to one. At a rest point every rate of change vanishes, so all strategies with positive frequency have equal fitness. We call a rest point *stable* when it is asymptotically stable, and we describe it by the strategies present, of which there may be one, two, three, or four (mono-, di-, tri- or four-morphic). A parametrization can have one, two or three stable rest points; when there are multiple, a population evolves to the one in whose basin of attraction it starts. If a parametrization has one stable rest point, we refer to it (and the corresponding division of labor) as monostable. If a parametrization has two (three) stable rest points, they are bistable (tristable).

For our results below we focus on the setting in which both tasks have identical payoff parameters (i.e.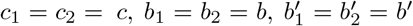), as our goal is to explore the effects of interactions rather than asymmetries between the tasks. Identical payoff parameters also allow us to derive the thresholds between all evolutionary outcomes. When the tasks have non-identical parameters, not all thresholds can be solved for analytically, and those that can be solved have more parameters and are harder to interpret.

With identical payoff parameters, both tasks are the same two-strategy game: anti-coordination, defined above; donation, in which *D* is the dominant strategy, defined by *b < c* and *b*^*′*^ *− c < b*; or coordination, in which players get higher payoff for taking the same action as each other (i.e. *b < c* and *b*^*′*^ *− c > b*). Our results below cover the first two settings. We report results for two interacting anti-coordination games in the main text, and report similar results with two donation games in Appendix C.

Evolutionary dynamics when the tasks are independent (i.e. *δ* = 0 and *γ* = 0) are detailed by Cressman et al. (2000) and Hashimoto (2014). Let 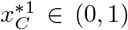 and 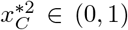 denote the evolutionarily stable frequencies of performers of tasks one and two, respectively, when they are analyzed separately. With no interactions between tasks, the set of stable rest points is a line of points *S* that sits at the intersection of the planes 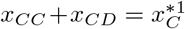 and 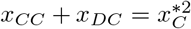 with the simplex *x*_*CC*_ + *x*_*CD*_ + *x*_*DC*_ + *x*_*DD*_ = 1 (Fig. 1b). Along *S*, the frequency of performers of each task equals its single-task equilibrium value, the two planes together imply 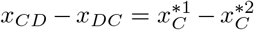, and no point of *S* is favored over another: a population that starts off *S* evolves to some point on it, and a perturbation moves it to another point on it. We ask how an interaction between the two tasks changes this outcome.

**Figure 1:**
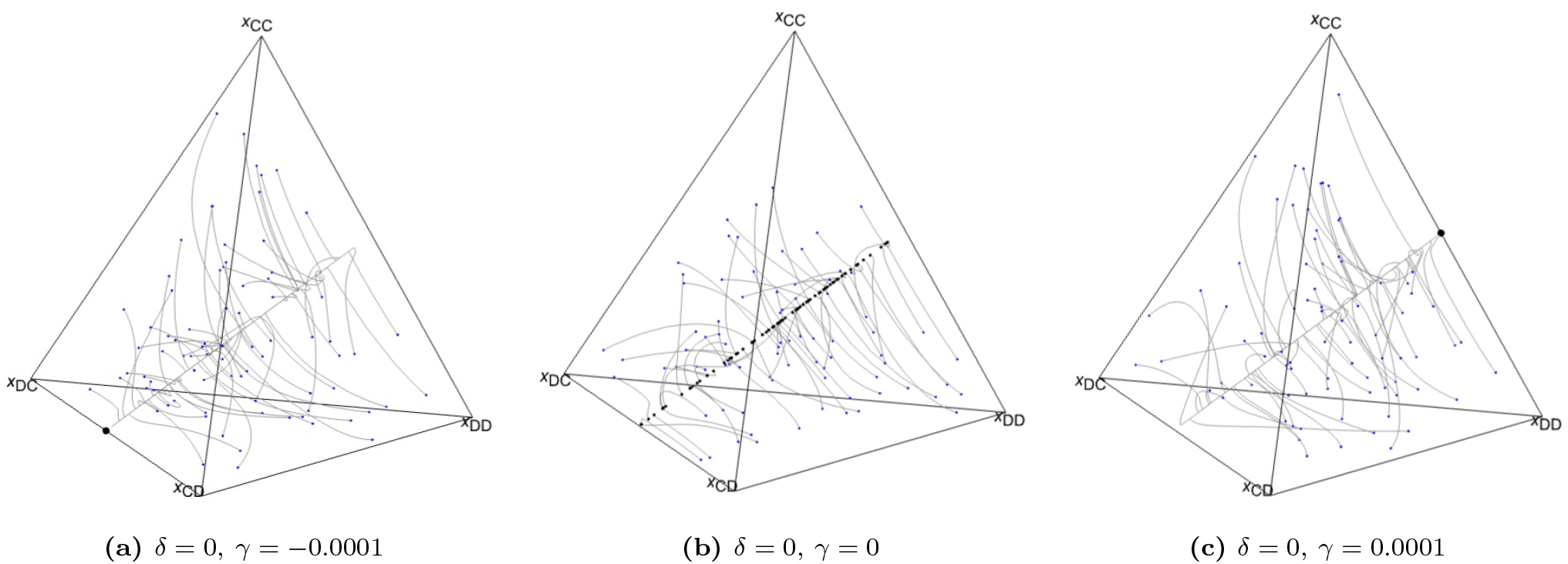
An arbitrarily small interaction between tasks replaces the line of rest points with isolated stable equilibria. In each subplot, a given point in or on the simplex corresponds to a frequency distribution across the four strategies in the population. Each corner is a population composed entirely of one strategy. Gray lines indicate evolutionary trajectories with randomly initialized frequencies of strategies, indicated by small blue dots. The large black dots in panels (*a*) and (*c*) indicate stable rest points. In the central panel (*b*) the tasks are independent, and the trajectories terminate along the line of rest points *S* rather than at a point. The endpoints of each trajectory are indicated by small black dots. An interaction term of magnitude 10^*−*4^, against payoffs of order unity, produces an isolated rest point: full specialization (*CD, DC*) in panel *a* and asymmetric generalism (*CC, DD*) in panel *c*. Parameters: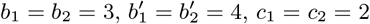.

## 3 Results

### 3.1 Any interaction between tasks replaces the line of rest points with isolated rest points

If there is any deviation from the additive payoff matrix, the line of rest points *S* of two independent anti-coordination games no longer exists. Adding an arbitrary deviation ϵ ∈ ℝ to any element of matrix (3) with *δ* = *γ* = 0 produces replicator dynamics that generically equilibrate at an isolated point. Complementary and antagonistic interactions between tasks are structured deviations of this kind, and they also produce isolated rest points (Fig. 1).

The magnitude of the interaction terms determines which composition is selected, but not whether the line is replaced by isolated points: interaction terms of magnitude 10^*−*4^ already produce isolated rest points (Fig. 1a,c). As the interaction terms are made smaller, the selected composition approaches an *endpoint* of the segment *S* rather than a point in its interior, and which endpoint depends on the ordering of *δ* and *γ* and, where two rest points are stable, on the initial composition. The line *S* is therefore a property of the additive model only, and not an approximation valid for weak interactions.

There is a simple reason why the additive model has a line of rest points. Along *S* all four strategies have equal fitness, which is why every point of the line is a rest point; the equality holds because the payoff an individual receives is the sum of two independent single-task payoffs, and the frequency of performers of each task is at its single-task equilibrium value. Adding a term to the payoff matrix that depends on the pair of strategies rather than on the two tasks separately removes that equality everywhere except at isolated compositions.

### 3.2 Properties of divisions of labor

When *δ* or *γ* is nonzero, the dynamics fall into twelve outcomes (Table 2; Figs. 2 and 3). Only outcome I (monomorphic double-cooperators) occurs with independent tasks. The divisions of labor in outcomes II–V are monostable when the tasks interact, but with independent tasks they are only neutrally stable endpoints of the line *S*. We also show in Appendix C that in the setting in which both tasks are donation games, dynamics can be classified into eight different outcomes, seven of which are not possible under the model of independent tasks.

**Table 2:** Equilibrium divisions of labor when both tasks are identical anti-coordination games. Sets of strategies in parentheses indicate a polymorphic division of labor containing those strategies. In outcomes with only one set (I–V and XII), the equilibrium division of labor is a single, globally stable point that contains the indicated strategies. In outcomes with an “or” between strategy sets, the two divisions of labor on either side of the “or” are multistable, and a population could evolve to either depending on its initial strategy distribution and the underlying parameters.

| Label | Equilibrium division(s) of labor | Description |
| --- | --- | --- |
| I. | $CC$ | Monomorphic double-cooperators |
| II. | $(CC, DD)$ | Asymmetric generalism |
| III. | $(CD, DC)$ | Full specialization |
| IV. | $(CC, CD, DC)$ | Partial (cooperator-)specialization |
| V. | $(DD, CD, DC)$ | Partial (defector-)specialization |
| VI. | $(DD, CD)$ or $(DD, DC)$ | Bistable dimorphisms with double-defectors and one specialist |
| VII. | $(CC, DD)$ or $(CD, DC)$ | Bistability between II and III |
| VIII. | $(CC, DD)$ or $(CC, CD, DC)$ | Bistability between II and IV |
| IX. | $(CC, DD)$ or $(DD, CD, DC)$ | Bistability between II and V |
| X. | $CC$ or $(DD, CD)$ or $(DD, DC)$ | Tristability between I and the two dimorphisms in VI |
| XI. | $(CC, DD)$ or $(DD, CD)$ or $(DD, DC)$ | Tristability between II and the two dimorphisms in VI |
| XII. | $(CC, CD, DC, DD)$ | Four-type division of labor with all strategies present |

**Figure 2:**
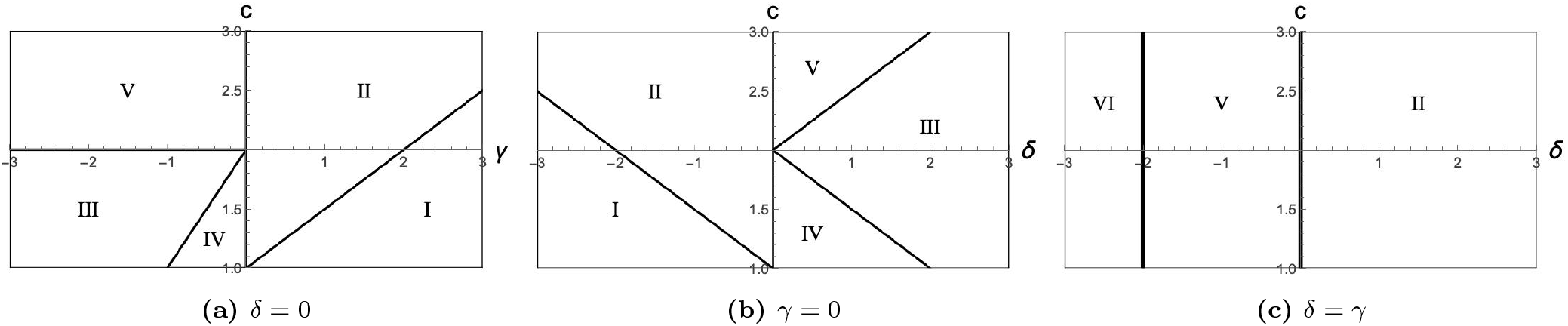
Complementarity for specialists and double-defectors promotes specialization more than antag-onism for double-cooperators does. Each Roman numeral corresponds to a distinct evolutionary outcome when both tasks are identical anti-coordination games (Table 2), and thick black lines are the boundaries between outcomes in parameter space. The interaction term for double-cooperators, *γ*, which includes both public and private interactions (panel a), and the interaction term for all other strategies, *δ*, which includes only public interactions (panel b), have nearly symmetric effects on the equilibrium division of labor. But full specialization (III) and cooperator-specialization (IV) occupy larger regions with complementarity for specialists and double-defectors only (*δ >* 0, *γ* = 0; panel b) than with antagonism for double-cooperators only (*γ <* 0, *δ* = 0; panel a). Panel c shows the case *δ* = *γ*. The region of (*c, δ*)-space corresponding to full specialization (III) in Fig. 2*b* is larger than the corresponding region of (*c, γ*)-space in Fig. 2*a* by an area of 1.5 within the plotted range; the region of (*c, δ*)-space corresponding to cooperator-specialization (IV) in Fig. 2*b* is larger than the corresponding region of (*c, γ*)-space in Fig. 2*a* by an area equal to one half. Parameters: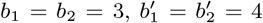, and *c*_1_ = *c*_2_ = *c* on the vertical axis.

**Figure 3:**
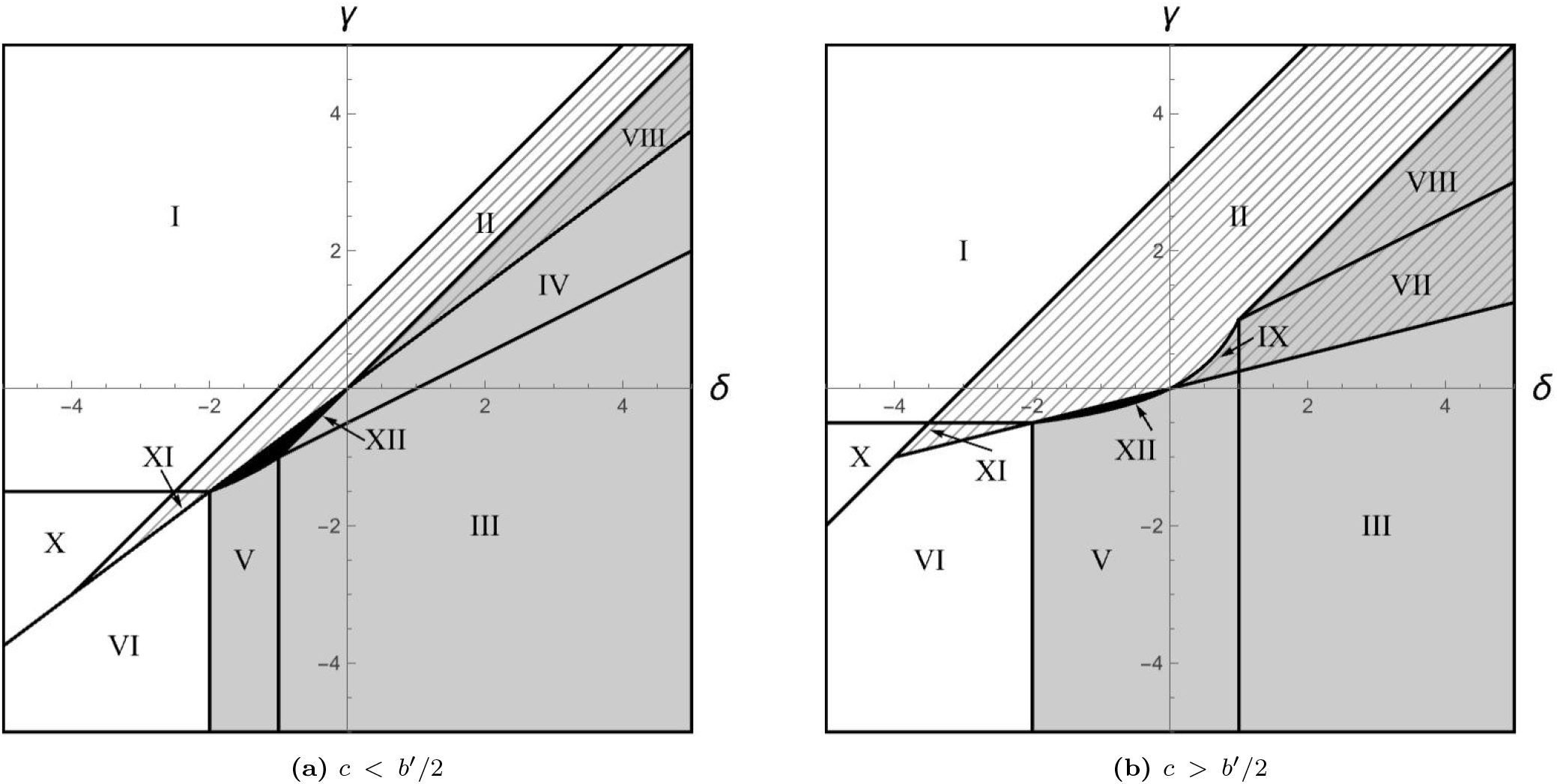
Interactions between two identical anti-coordination tasks produce twelve evolutionary outcomes, including bistability between specialized and asymmetric divisions of labor. Each Roman numeral corresponds to a distinct evolutionary outcome when both tasks are identical anti-coordination games (Table 2). Several outcomes entail two or three divisions of labor that are potential evolutionary equilibria, depending on the initial strategy distribution. In the gray regions, (one of) the equilibrium divisions of labor contains a positive frequency of both specialists (that is, *CD* in task one and *DC* in task two). In the striped regions, (one of) the equilibrium divisions of labor is the asymmetric generalist equilibrium (*CC, DD*). In the three regions (VII, VIII, and IX) that are both gray and striped, one equilibrium division of labor is (*CC, DD*), and the other contains positive frequencies of both specialists. In each of these bistable outcomes, individuals in the equilibrium division of labor containing both specialists have higher fitness than individuals in the asymmetric generalist equilibrium. In the black region (XII), the evolutionary outcome is a globally stable division of labor with positive frequencies of all four types. Parameters: *b*_1_ = *b*_2_ = 3 and 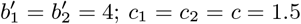 in panel a and *c* = 2.5 in panel b.

Six of the outcomes are monostable: one monomorphic, two dimorphic, two trimorphic, and one four-morphic. When the interaction term for double-cooperators, *γ*, is large enough, the population consists only of double-cooperators (I). The other monostable outcomes (II–V) occur at intermediate values of (*δ, γ*). In asymmetric generalism (II), double-cooperators perform all tasks and double-defectors perform none; in full specialization (III), the two specialists coexist, at equal frequency when the tasks are identical. In the two trimorphisms, partial cooperator-specialization (IV) and partial defector-specialization (V), double-cooperators or double-defectors co-exist with both specialists, which typically have much higher frequency. The four-morphic division of labor (XII) occurs where several regions of stability overlap, and it is the only outcome in which all four strategies coexist.

Four of the twelve outcomes entail bistable divisions of labor. In three of these (VII–IX), the asymmetric generalist equilibrium is bistable with one of the fully or partially specialized equilibria, and individuals in the specialized division of labor have higher fitness (Section 3.5). In the remaining bistable outcome (VI), the two equilibria contain the double-defectors and either specialist type (but not both). These equilibria occur because the antagonism between tasks experienced by specialists (*δ <* 0) is strong enough that either specialist type has higher fitness if the other is absent. Two of the outcomes entail tristable divisions of labor (X, XI). In these two outcomes, the two dimorphisms of outcome VI (double-defectors coexisting with one specialist type) are tristable with either the monomorphic cooperator equilibrium or the asymmetric generalist equilibrium. While the one-specialist divisions of labor (VI) have equal fitness if the tasks have identical payoff parameters, they can have higher or lower fitness than the monomorphic cooperator equilibrium in X or the asymmetric generalist equilibrium in XI depending on parameter values.

### 3.3 Bifurcation thresholds between outcomes

We have derived the thresholds between evolutionary outcomes as functions of the interaction terms *δ* and *γ* (Figs. 2 and 3). We report them here for identical tasks 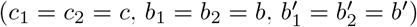.

The thresholds between outcomes clarify the strategic incentives that produce them. In a few cases, the explanation depends directly on how the task interactions change the ordering of the payoffs in (3). For example, in region I in Figs. 2 and 3, double-cooperation (*CC* ) is the dominant strategy among all four, and the corresponding equilibrium is monomorphic double-cooperators. A second example is the vertical threshold *δ* = *b*^*′*^ *−* 2*b* between regions V and VI (Fig. 3). We call the two-strategy game played between a pair of strategies, here the specialists *CD* and *DC*, a constituent game. The constituent game between the two specialists is an anti-coordination game when *δ > b*^*′*^ *−* 2*b*, so that each specialist has higher fitness when the other is present, and a coordination game when *δ < b*^*′*^ *−* 2*b*, so that each has higher fitness when the other is absent. The constituent game between double-defectors and either specialist does not contain the interaction term and remains an anti-coordination game. This change in the constituent game between the specialists explains why the equilibrium changes from monostable (*DD, CD, DC*) in region V to bistable (*DD, CD*) or (*DD, DC*) in region VI.

All of the bifurcation thresholds discussed in Sections 3.3.1–3.3.3 below are located in regions of parameter space in which the constituent games played by all pairs of existing strategies at equilibrium are anti-coordination games. Neither dominance nor a change in a constituent game explains these thresholds, and in each of these regions the equilibrium is monostable asymmetric generalism, monostable partial or full specialization, or bistability between the two.

#### 3.3.1 Interpreting the thresholds between partially- and fully-specialized divisions of labor

Consider the vertical (*δ*) threshold between regions V (partial defector-specialization, (*DD, DC, CD*)) and III (full specialization, (*CD, DC*)) (Fig. 3a,b), which is *δ* = *−b*^*′*^ + 2*c*. Above this threshold full specialization can be an equilibrium and partial defector-specialization cannot, and below it the reverse. In a population where only double-defectors and both specialist types are present, increasing *b*^*′*^ increases the fitness of both specialists, who have a chance of encountering their own type and receiving *b*^*′*^ in task benefits. Conversely, increasing *c* decreases the fitness of both specialists, because they always pay cost *c* to perform one task. Changing either *b*^*′*^ or *c* does not change the fitness of double-defectors. Thus, the public task interaction effect *δ* received by specialist types when they encounter each other can compensate for the net effects of *b*^*′*^ (which benefits specialists) and *c* (which costs specialists) on fitness. Consider a population in division of labor V, with *δ < −b*^*′*^ + 2*c*. As *δ* increases, the fitness of both specialists increases while that of double-defectors does not change, and *x*_*DD*_ decreases relative to *x*_*CD*_ = *x*_*DC*_. As *δ* approaches *−b*^*′*^ + 2*c* from below, *x*_*DD*_ approaches 0. When *δ* = *−b*^*′*^ + 2*c*, the stable rest points III and V coincide.^*^

The threshold between regions III and IV (partial cooperator-specialization, (*CC, CD, DC*)) in Fig. 3a is 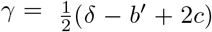. Above it, double-cooperators coexist with both specialists and partial cooperator-specialization is the stable rest point; below it, full specialization is. In a population where only double-cooperators and both specialists are present, specialists have a payoff advantage when they encounter double-cooperators, which is equal to *π*(*CD*|*CC*) *− π*(*CC*|*CD*) = *δ − γ* + *c*; meanwhile, double-cooperators have an advantage when they encounter their own type, equal to *π*(*CC*|*CC*) *− π*(*CD*|*CD*) = *γ* + *b*^*′*^ *− c*. The threshold is the value of *γ* at which these two advantages are equal.

#### 3.3.2 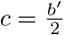 is a boundary between two different spaces of possible divisions of labor

The arrangement of outcomes in the bifurcation diagram depends on the orientations in (*δ, γ*)-space of the two thresholds described in Section 3.3.1 (see Fig. 3*a* versus 3*b*). The thresholds intersect at the point (*δ, γ*) = (*−b*^*′*^ +2*c, −b*^*′*^ +2*c*). When 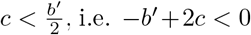, the thresholds intersect in the lower left quadrant of (*δ, γ*)-space, and the bifurcation diagram is isomorphic to Fig. 3*a*. When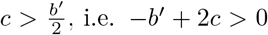, the thresholds intersect in the upper right quadrant, and the bifurcation diagram is isomorphic to Fig. 3*b*. This change occurs because the vertical crossing of these two thresholds as *c* increases from *c < b*^*′*^*/*2 to *c > b*^*′*^*/*2 makes the parameter regions in which defector-specialization (*DD, DC, CD*) and full specialization (*CD, DC*) are stable *overlap* with the region in which asymmetric generalism (*CC, DD*) is stable, creating two new bistable outcomes and removing monostable cooperator-specialization (IV). When 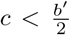, the threshold 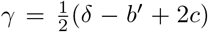 between monostable cooperator-specialization (IV) and full specialization (III) is *below* the threshold *γ* = *δ*(*b − c*)*/*(2*b − b*^*′*^) between region IV and region VIII (Fig. 3*a*), which is the lower threshold above which asymmetric generalism (*CC, DD*) can be a stable rest point. When 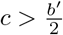, the line 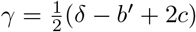 is now the threshold between regions VII and VIII, *above* the line *γ* = *δ*(*b − c*)*/*(2*b − b*^*′*^), which is now the threshold between regions III and VII. Thus, the region in which full specialization is a stable rest point has increased in height and now overlaps with the region in which asymmetric generalism is a stable rest point, creating a new bistable evolutionary outcome (VII).

#### 3.3.3 An efficiency requirement δ > γ is necessary for specialists to persist in five of the seven outcomes that contain them

The threshold *δ* = *γ* between regions II (monostable asymmetric generalism, (*CC, DD*)) and VIII (bistability between (*CC, DD*) and (*CC, CD, DC*)) has special importance. First, *γ < δ* is necessary for either full specialization (*CD, DC*) or cooperator-specialization (*CC, CD, DC*) to be equilibria. Second, if *δ >* 0, then *δ > γ* is a requirement for defector-specialization (*DD, CD, DC*) to be an equilibrium as well as full- and cooperator-specialization^*†*^, and these are the only equilibria that contain both specialist types when *δ >* 0. These two requirements are implied by the upper boundaries of regions III, IV, and VIII in Fig. 3*a*, and those of regions III, V, VII, VIII, and IX in Fig. 3*b*. The condition *δ > γ* can be interpreted as an efficiency requirement: specialists must obtain more from the interaction between tasks than double-cooperators do. In other words, there is a negative private interaction between tasks, e.g., due to the increased cost of having to allocate resources to both.

The threshold *δ* = *γ* can also be interpreted as the point at which the opposing impacts of the task interaction on the fitness of double-cooperators and specialists cancel out. Double-cooperators receive *γ* in every encounter, whereas specialists receive *δ* only in encounters with a double-cooperator or with the other specialist type. Therefore if 0 < *δ* ≤ *γ*, double-cooperators receive a complementarity term in every encounter that is at least as large as the complementarity term that specialists receive in a fraction of their encounters, and either monostable asymmetric generalism (II) or monomorphic cooperation (I) is the evolutionary outcome. By contrast, if *δ > γ*, specialists compensate for the decreased frequency with which they receive the task-interaction effect by having a higher task-interaction effect each time they receive it. This compensation allows the pair of specialist strategies to coexist with double-cooperators or even outcompete them when 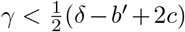 (see Section 3.3.1 above). The line *δ* = *γ* is therefore a special case with only three evolutionary outcomes: asymmetric generalism (II) when the tasks are complementary (*δ* = *γ >* 0) and defector-specialization or bistability between defectors and either specialist (VI) when they are antagonistic (*δ* = *γ <* 0; see Fig. 2*c*).

### 3.4 Task complementarity for specialists and double-defectors (δ > 0) promotes specialization more than antagonism for double-cooperators (*γ <* 0) does

When one interaction term is zero, the effects of *δ* and *γ* are nearly symmetric about the vertical axis, but full specialization (III) and cooperator-specialization (IV) occupy larger regions when *δ >* 0 (Fig. 2b) than when *γ <* 0 (Fig. 2a), and defector-specialization (V) occupies a correspondingly larger region when *γ <* 0. The thresholds of Section 3.3.1 explain this asymmetry. The threshold *δ* = *−b*^*′*^ + 2*c* between regions III and V does not depend on *γ* (Fig. 2a), because *γ* only affects double-cooperators, while *δ* only affects specialists in outcomes III and V. As *δ* increases above 0 (with *γ* = 0), the minimum cost *c* that is required for (*DD, CD, DC*) to be an equilibrium instead of (*CD, DC*) increases (∂*c/*∂*δ* = 1*/*2; Fig. 2*b*), enlarging the region of parameter space in which full specialization occurs relative to the upper left quadrant in Fig. 2*a*. The threshold 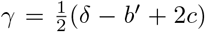 between regions III and IV increases faster with *γ* (∂*c/*∂*γ* = 1) than it decreases with *δ* (∂*c/*∂*δ* = *−*1*/*2). Thus antagonism for double-cooperators promotes cooperator-specialization only when |*γ*| is small, whereas complementarity for specialists and double-defectors promotes it up to large |*δ*|.

### 3.5 When specialization is bistable with asymmetric generalism, the division of labor with both specialists has higher fitness

In outcomes VII, VIII and IX (Fig. 3), asymmetric generalism (*CC, DD*) is bistable with full (*CD, DC*), cooperator- (*CC, CD, DC*), or defector-specialization (*DD, CD, DC*), respectively. We have proved (Appendix B) that individuals in the division of labor containing both specialists have higher fitness than individuals in the asymmetric generalist division of labor in these three outcomes.

In these outcomes, the tasks have a complementary interaction for all strategies, but the synergy benefit is greater for specialists and double-defectors than for double-cooperators (efficiency requirement *δ > γ* from Section 3.3.3). The requirement suggests a simple underlying reason that the divisions of labor containing both specialists have higher fitness than their asymmetric generalist counterparts: because specialists and double-defectors receive a larger complementarity term than double-cooperators, a population in which both specialists are present will have higher fitness at equilibrium than a population in which specialists are absent, as in asymmetric generalism.

### 3.6 Divisions of labor can exhibit hysteresis as task complementarity parameters vary

In the upper right quadrants of Fig. 3*a* and *b*, the parameter regions corresponding to bistable outcomes are in the middle, in the overlap between the stability threshold for asymmetric generalism and those of the specialized divisions of labor, while the regions corresponding to monostable outcomes are on the top (all cooperators or asymmetric generalism) and bottom (full or defector-specialization). If either interaction parameter fluctuates over a large enough range while the other remains constant, the division of labor therefore exhibits hysteresis (Fig. 4): whenever the parameter crosses a threshold, the equilibrium before the crossing is the initial condition afterwards, and the population evolves to the stable rest point whose basin of attraction contains the prior equilibrium.

**Figure 4:**
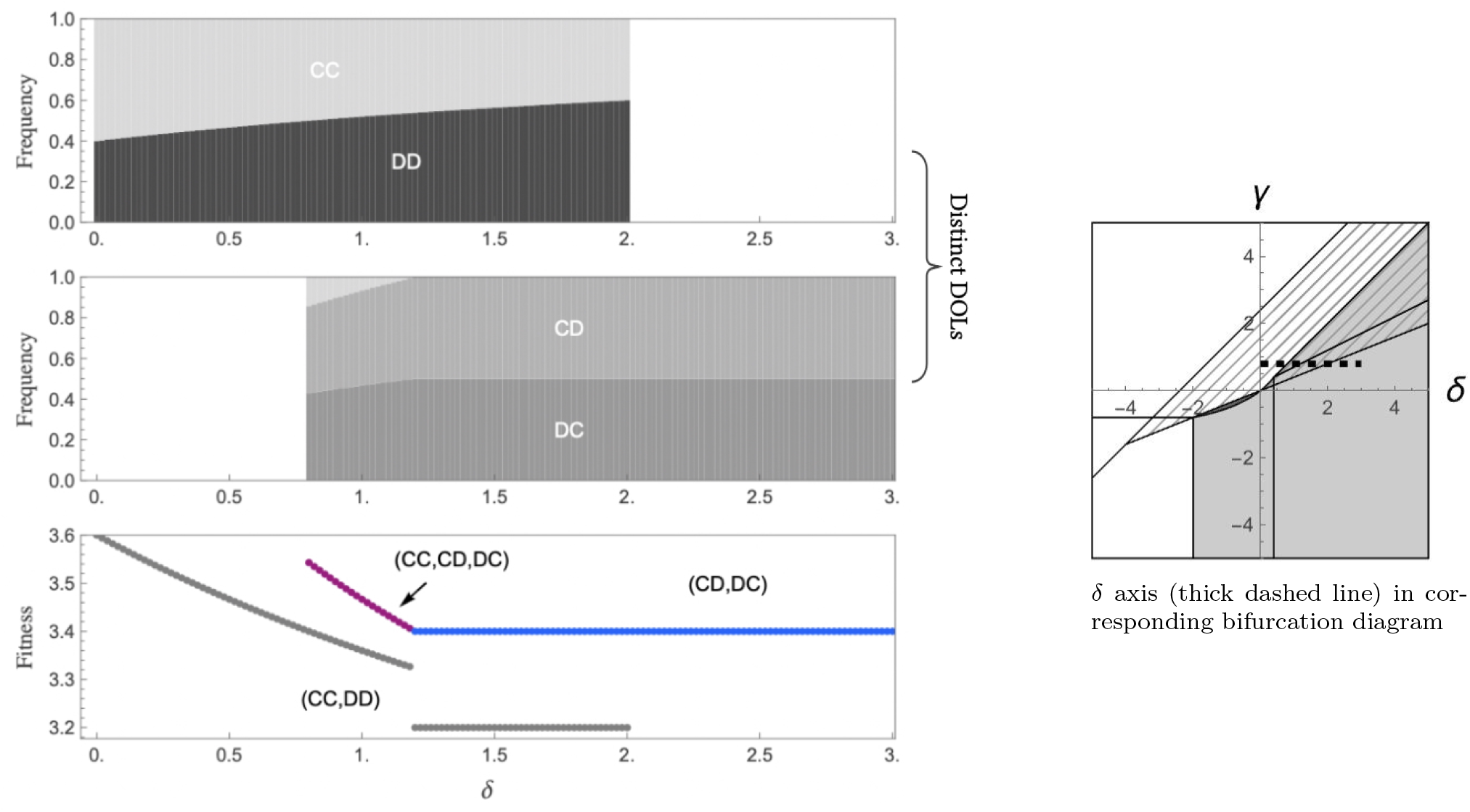
Bistable divisions of labor can exhibit hysteresis as *δ* varies, with consequences for fitness. The top two plots on the left represent distinct evolutionarily stable divisions of labor as a function of *δ*. The stacked bars in either plot at a given value of *δ* represent the frequencies of existing strategies at that equilibrium (see white labels). In either plot, the white space indicates that the division of labor is monostable and is shown in the other plot. Where both plots have stacked bars, the two divisions of labor are bistable. The lower plot shows the fitness of individuals at each division of labor; the specialized divisions of labor have higher fitness than asymmetric generalism. The right plot shows the transect along *δ* (thick dashed line) in the bifurcation diagram for these parameters, which is isomorphic to Fig. 3b. Parameters: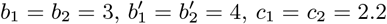, and *γ* = 0.8.

If an interaction parameter oscillates over time, the population alternates between long periods of asymmetric generalism and long periods of specialization (Fig. 4, top two panels). Because the specialized division of labor has the higher fitness in every outcome in which it is bistable with asymmetric generalism (Section 3.5), the population’s fitness exhibits hysteresis as well as its division of labor (Fig. 4, bottom panel).

## 4 Discussion

We have studied how interactions between two tasks shape the evolution of the division of labor, in a model in which performing both tasks changes the payoff by one amount for individuals who perform both and by another amount for individuals who perform at most one. That single assumption produces the large space of evolutionary outcomes described above. Task interactions permit the evolution of specialized divisions of labor, and in most cases this requires that the interaction effect be more favorable for specialists and double-defectors than for double-cooperators (*δ > γ*). When both tasks are identical anti-coordination games and a specialized division of labor is bistable with asymmetric generalism, the specialized division of labor always has higher fitness. Bistability also implies that a population exhibits hysteresis when an interaction parameter fluctuates, alternating between specialized and non-specialized divisions of labor with correspondingly higher and lower fitness.

Our results can inform predictions about how the division of labor on two tasks evolves in natural populations. It is difficult to identify the correct game to model a given task in a natural population; for example, when a human forager collects vegetation and shares it with another, how many net calories are gained and what variety of nutrients is provided? By contrast, it can be easier to identify if two tasks are complementary or antagonistic when performed together and if the interaction is more or less favorable to specialists than generalists. In our example, if humans forage both vegetation and meat, they ingest a wider variety of nutrients than if only one of these foods is foraged, and the nutrients from both can have a synergistic effect on payoff. Moreover, an individual who hunts and gathers on the same day divides a fixed budget of time and energy between the two activities, whereas two individuals who divide the activities between them do not, allowing specialists to extract more synergistic benefits from the tasks than double-cooperators do (Bird, 1999; Hooper et al., 2015). A similar efficiency advantage for specialists appears in colonies of *P. aeruginosa*: production of pyoverdine (task one) suppresses that of pyochelin (task two), so a cell that produces both siderophores obtains less from the pair than two cells that divide them (Kramer et al., 2020; Ross-Gillespie et al., 2015).

If either complementarity or antagonism is observed between the tasks a species performs, and this effect may differ between performers of both tasks and performers of at most one, our results imply several general predictions about the species. Under the hypothesis that the tasks are either both anti-coordination or both donation games, it is likely that one or both of the asymmetric generalist (*CC,DD* ) and fully- or partially specialized (*·,CD,DC* ) divisions of labor are stable equilibria. Further, if the tasks are anti-coordination games and both asymmetric generalism and specialization are observed in similar populations of the same species, the specialized population likely obtains more payoff from the tasks in the long run, because the task interaction is more favorable to specialists and double-defectors than to double-cooperators.

The condition for specialization, that performers of at most one task receive a more favorable interaction effect than performers of both, corresponds to the standing requirement in the theory of division of labor that specialists must be more efficient than generalists at the task they perform (Cooper et al., 2021; Cooper and West, 2018; Rueffler et al., 2012). In those models the efficiency benefit is a property of a single individual performing a single task, expressed as an accelerating performance or a convex trade-off. The difference *γ − δ*, the amount an individual gains or loses by covering both tasks itself rather than leaving one to a coplayer, is the counterpart of that benefit in a model where the tasks are games rather than allocations of effort. If this difference is negative, an individual covering both tasks does worse than specialists covering one each; this is the counterpart of the efficiency requirement. Our results indicate that the requirement still holds when the two tasks interact, and that complementarity between the tasks *per se* does not substitute for it.

Several assumptions of the model are worth highlighting. The most consequential is that the interaction between tasks is realized within an interacting pair, so that a specialist collects its interaction term only when its coplayer performs the task it omits. Realizing the interaction within a pair creates an asymmetry between an individual who covers both tasks and one who does not, and our results depend on that asymmetry. A model in which the interaction instead depends on the total amount of each product present in the population (or a larger neighborhood) need not display the same level of asymmetry. Freely diffusing products such as siderophores or plant volatiles are the case where the alternative formulation is the more natural one, and we have not analyzed it. A second assumption is that individuals are paired uniformly at random, so there is no assortment between strategies. Because the interaction between tasks is realized by pairs, we expect assortment to matter here more than it does in a single game, and in particular we expect positive assortment among specialists to widen the range of parameters over which they persist. Also, strategies are the four combinations of two binary actions, rather than continuous levels of effort on each task; a continuous model would let effort on the two tasks diverge gradually and would replace our discrete penalty *γ − δ* with the curvature of an allocation trade-off, which is the form the efficiency requirement takes in the continuous division-of-labor literature (Rueffler et al., 2012). The two interaction terms are constants, whereas in a metabolic or a time-budget setting the penalty for performing both tasks plausibly depends on how much of each task is performed, and hence on the composition of the population. Finally, the games are one-shot and unconditional, so none of the linkage between tasks that arises from conditioning behavior in one game on behavior in the other is present (Donahue et al., 2020; Salahshour, 2022). The purpose of a model this simple is not to give a quantitative account of siderophore production or of foraging in any particular population, but to identify which features of an interaction between two tasks determine how the labor of performing them is distributed.

Several extensions of the model remain to be analyzed. First, the evolutionary outcomes of task interactions have not been described for five further pairs of two-strategy games: two coordination games; anti-coordination and donation games; coordination and donation games; coordination and anti-coordination games; and donation and harmony games, the last of which has cooperation as a dominant strategy. Second, an additional parameter could separate the interaction effect on specialists from that on double-defectors, which would show whether our outcomes depend on the two receiving the same interaction term. Third, task-performance encounters could include more than two individuals; we expect our results to extend to larger groups only under conditions that have yet to be established. Fourth, the division of labor could involve three or more tasks, which greatly increases the number of possible outcomes. Finally, if two bistable divisions of labor produced by task interactions are both present on patches of a metapopulation with uniform dispersal, the division of labor to which the population will evolve is not known.

## Supporting information

Electronic supplementary material

## Appendix

### A Stability analysis of rest points

Following Section 2, the matrix (3) gives the total payoff from both tasks that each of the four strategies (*CC, CD, DC, DD*) receives when they play the task-game against any other strategy. The replicator dynamics of strategy frequencies are at a rest point if *∀g ∈ {CC, CD, DC, DD}*,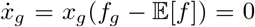. There are fifteen symbolic solutions to this system of equations, which are candidate equilibria for a specific parametrization. Because the frequencies sum to one, we write *x*_*DD*_ = 1 *− x*_*CC*_ *− x*_*CD*_ *− x*_*DC*_ and denote the vector-valued function giving the rates of change of the three remaining frequencies as 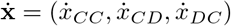, and denote the frequency distribution at a rest point as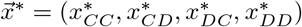, where “*” indicates the rest point value.

Rest point 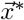 is asymptotically stable if any population whose frequency distribution starts in the basin of attraction of 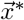 evolves to and equilibrates at 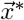; that is, 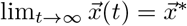 for every initial frequency distribution 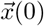 in the basin of attraction of 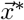. A sufficient condition for 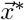 to be asymptotically stable is that every eigenvalue of the Jacobian matrix of 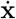 evaluated at 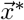 has negative real part. If some eigenvalues have zero real part and the others negative real part, asymptotic stability of 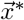is possible but not certain, and further analysis is required to determine stability. The Jacobian matrix **J** is written

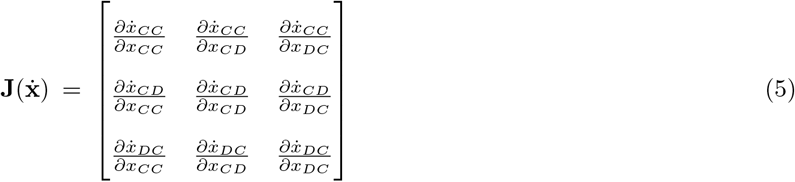

To derive the region of parameter space in which a given rest point 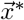 is asymptotically stable, we compute the eigenvalues of **J** evaluated at 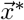 and simplify the conditions for all eigenvalues to have negative real part. This computation allows us to solve for the bifurcation thresholds between regions corresponding to different equilibria as functions of parameters.

If no candidate equilibrium has all eigenvalues with negative real part, but one or more candidates have eigen-values with zero real part and none with positive real part, we can determine whether the latter are asymptotically stable by numerically integrating the replicator dynamics until equilibration for a large sample of trajectories initialized randomly within the simplex. In the case of two independent tasks (*δ* = *γ* = 0), the Jacobian at every point along the line of rest points *S* has a zero eigenvalue, corresponding to the direction along *S*. Given an initial frequency distribution 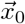, we can compute the equilibrium along this line by numerically integrating the replicator dynamics; a large sample of these trajectories generates the representation of *S* shown in Fig. 1*b*.

The numerical integrator that we used was Mathematica’s NDSolve (see electronic supplementary material). In Fig. 1, each panel shows a collection of 200 trajectories, each with initial frequencies that were initialized uniformly at random such that *∀q ∈ {CC, CD, DC, DD}, x*_*q*_ *∈* [0, 1], and 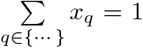 . For each trajectory, the replicator ODEs were integrated until stoptime *t*_max_ = 5 *×* 10^6^. For the parametrizations in Fig. 1, strategy frequencies evolve to and stabilize at their long-run values (i.e. those at *t*_max_) at an earlier timepoint than *t* = 10^6^. Thus, we are able to define the equilibrium of a given trajectory as the vector of strategy frequencies present at *t*_max_. Because of the simplicity of this system, initializing and integrating two-hundred trajectories until *t*_max_ takes a fraction of a second.

The bifurcation thresholds corresponding to each evolutionary outcome in Fig. 3*b* are shown in Fig. 5. The thresholds in Fig. 3*a* (with 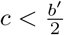) are isomorphic to those in 3*b* (with 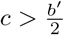), with one exception: in Fig. 3*a*, the threshold between regions III and IV is 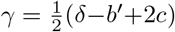 and that between regions IV and VIII is *γ* = *δ*(*b−c*)*/*(2*b−b*^*′*^).

**Figure 5:**
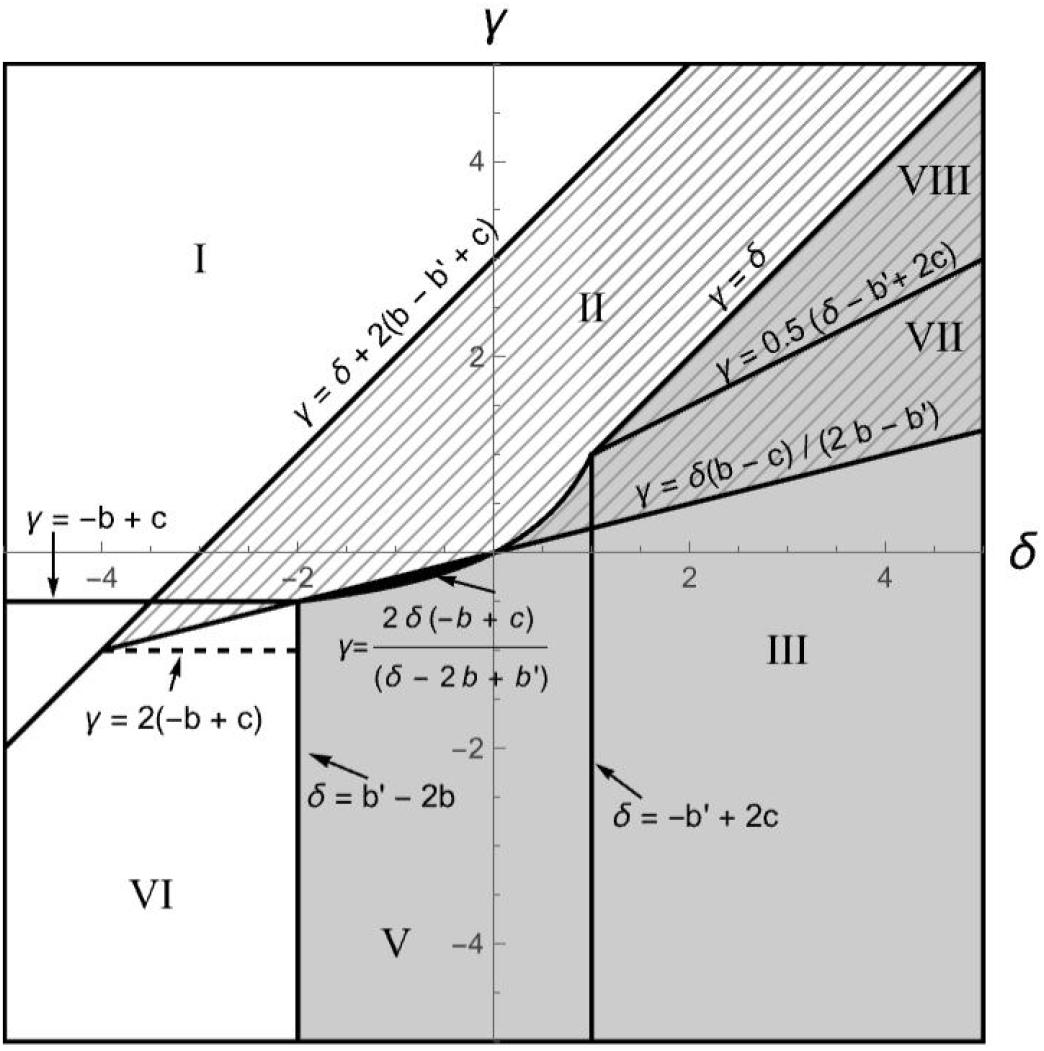
Bifurcation thresholds between evolutionary outcomes in (*δ, γ*)-space. Parameters as in Fig. 3b. For visual clarity, regions IX–XII are unlabeled, but can be found by referring to Fig. 3*b*.

As described in Section 3.3.2, these two thresholds exchange their vertical order in (*δ, γ*)-space when *c* increases above 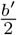, resulting in a distinct bifurcation diagram.

### B Derivation of bifurcation thresholds and fitnesses of equilibria

Derivation of thresholds and fitnesses of bistable equilibria can be performed using sections 1.1–1.3 of the Mathematica notebook in the electronic supplementary material. In section 1.1, we derive the symbolic expressions for the frequencies of strategies at each distinct rest point. By plugging these expressions for a given rest point into the expected payoff function, we derive the expressions for the fitness of individuals in a population whose division of labor is at that rest point. In section 1.2, we determine the bifurcation thresholds and regions of multistability between equilibria by algebraically simplifying the condition that all eigenvalues of the Jacobian evaluated at a given rest point are negative, using Mathematica’s Reduce function. When this condition is met, the given rest point is asymptotically stable; when the condition is met simultaneously for two or more rest points, those points are multistable.

In section 1.3, we prove that if both tasks are identical anti-coordination games, the specialized rest points have higher fitness than the asymmetric generalist rest points in outcomes VII, VIII and IX. An outline of this simple proof by contradiction is as follows. For any pair of bistable rest points from these three outcomes, say (*CD, DC*) and (*CC, DD*) in outcome VII, we assume the mean fitness of the specialized population is less than or equal to the mean fitness of the generalist population. Then, we show that all eigenvalues of the Jacobian matrices (5) evaluated at the respective rest points cannot simultaneously be negative. In other words, if the fitness at the specialized rest point is lower than or equal to the fitness at the asymmetric generalist rest point, the two rest points cannot both be stable; therefore, in the given outcome, the specialized rest point must have higher fitness than its alternative. Here we provide some analytical detail on the welfare-analysis proof corresponding to outcome VII, in which the fully specialized population (*CD, DC*) has higher fitness than the asymmetric generalist population (*CC, DD*).

The symbolic expression for the difference in fitness between the (*CC, DD*) and (*CD, DC*) rest points is

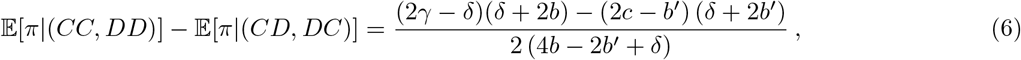

where E[*π*|(*CC, DD*)] is the fitness of individuals in the asymmetric generalist equilibrium and E[*π*|(*CD, DC*)] is the fitness of individuals in the fully specialized equilibrium. By simplifying the conditions for the appropriate eigenvalues to be simultaneously negative (see electronic supplementary material section 1.2), we have derived the following conditions for the rest points (*CD, DC*) and (*CC, DD*) to be bistable when both tasks are identical anti-coordination games:

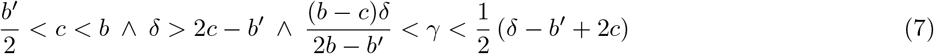

The denominator 2(4*b −* 2*b*^*′*^ + *δ*) of the fitness difference (6) is positive, by the first two inequality conditions in (7). Also, 2*γ − δ <* 2*c − b*^*′*^, by the rightmost inequality 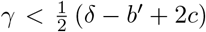 in (7). The first two inequalities in (7) give 2*c − b*^*′*^ *>* 0 and *δ >* 0, and 0 *< b < b*^*′*^ by the conditions for an anti-coordination game. Therefore (2*γ − δ*)(*δ* + 2*b*) *<* (2*c − b*^*′*^)(*δ* + 2*b*) *<* (2*c − b*^*′*^) (*δ* + 2*b*^*′*^), so the numerator in (6) is negative, which implies that the fitness difference (6) is negative when conditions (7) are met. In conclusion, in order for the fully specialized (*CD, DC*) and asymmetric generalist (*CC, DD*) equilibria to be bistable as in outcome VII, it is necessary that individuals in the asymmetric generalist equilibrium have lower fitness than their fully specialized counterparts.

### C Analysis of two identical interacting donation games

We applied the same analysis to two identical interacting donation games, in which performing a task alone costs more than it returns (*b < c*). In the absence of interactions between tasks, *DD* is the dominant strategy, and fixation of *DD* is the equilibrium of all evolutionary trajectories.

When the tasks interact, there are seven additional evolutionary outcomes that can occur, which are summarized in Table 3. The parameter regions for these outcomes are shown in Fig. 6, along with the functions for the thresholds between these regions. Note that outcome XV (bistability between cooperator-specialization and monomorphic defection) is not pictured; that outcome occurs when the threshold *γ* = 2(*−b*+*c*) exceeds the point *γ* = *δ* = *−b*^*′*^ +2*c*, which creates a new zone of overlap between stability regions for the monomorphic defection and cooperator-specialization divisions of labor. The key threshold value at which this change happens is 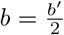.

**Table 3:** Equilibrium divisions of labor when both tasks are identical donation games. Sets of strategies in parentheses indicate a polymorphic division of labor containing those strategies. In outcomes with only one set (I, II, and XVI), the equilibrium division of labor is a single, globally stable point that contains the indicated strategies. In outcomes with an “or” between strategy sets, the two divisions of labor on either side of the “or” are multistable, and a population could evolve to either depending on its initial strategy distribution and the underlying parameters.

| Label | Equilibrium division(s) of labor | Description |
| --- | --- | --- |
| I. | $CC$ | Monomorphic double-cooperators |
| II. | $(CC, DD)$ | Asymmetric generalism |
| VII. | $(CC, DD)$ or $(CD, DC)$ | Bistability between II and specialization |
| VIII. | $(CC, DD)$ or $(CC, CD, DC)$ | Bistability between II and cooperator-specialization |
| XIII. | $CC$ or $DD$ | Bistability between monomorphic cooperation or defection |
| XIV. | $(CD, DC)$ or $DD$ | Bistability between specialization and defection |
| XV. | $(CC, CD, DC)$ or $DD$ | Bistability between cooperator-specialization and defection |
| XVI. | $DD$ | Monomorphic defection |

**Figure 6:**
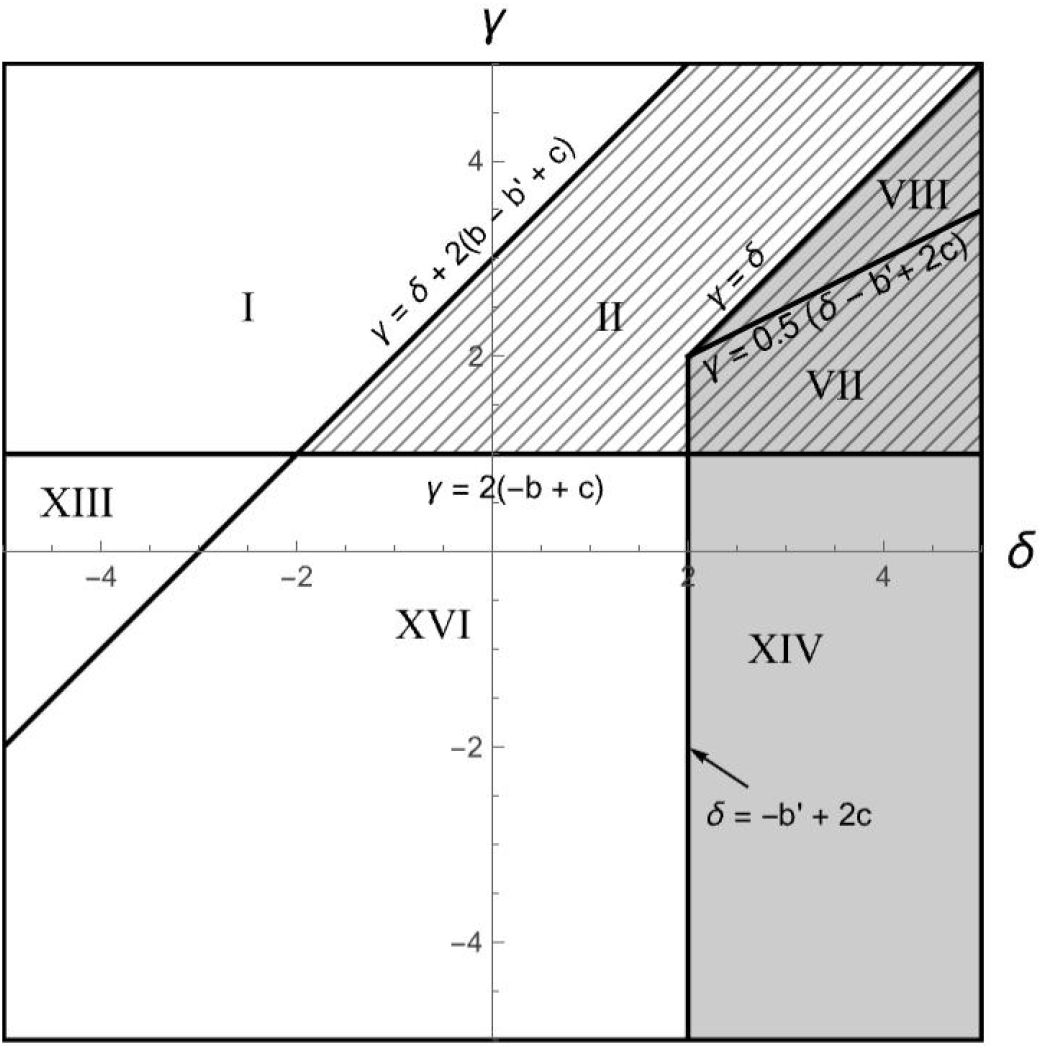
Evolutionarily stable divisions of labor on two identical tasks as a function of task interactions. Each Roman numeral corresponds to a distinct evolutionary outcome when both tasks are identical donation games (Table 3). In gray regions, one equilibrium division of labor contains both specialists; in striped regions, one equilibrium is asymmetric generalism (*CC, DD*). Parameters: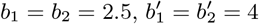, and *c*_1_ = *c*_2_ = *c* = 3.

For the bistable outcomes (VII, VIII, XIII, XIV, and XV), we compared the fitness of individuals at the alternative divisions of labor, as in Section 3.5. Using the proof by contradiction outlined in Appendix B, we prove that in outcomes XIV and XV individuals in the partially- or fully-specialized equilibrium have higher fitness than individuals in the monomorphic defector equilibrium with the same parameters. For the other bistable outcomes in this setting, the equilibrium division of labor with higher fitness depends on parameter values.

## Footnotes

* When double-defectors and specialists encounter each other, both receive the same benefit *b* from the one task performed, so this threshold does not depend on *b*.

† The exceptions are defector-specialization (V) and the four-type division of labor (XII), each of which can be the equilibrium when *δ <* 0 and *γ > δ* (Fig. 3*a*).

